# Otolith shape variability of labrid fish from Rapa Nui (Easter Island), southeastern Pacific

**DOI:** 10.1101/2024.04.05.588230

**Authors:** Andrés Castro-García, Erwan Delrieu-Trottin, Pablo Saenz-Agudelo, Cristian Rapu-Edmunds, Guido Plaza, Federico Márquez, Mauricio F. Landaeta

## Abstract

Fish otolith shape provides valuable insights into the taxonomic, phylogenetic, and ecological traits of fish species. This study aimed to assess and compare the phenotypic variation in otolith shape among four labrid species (*Anampses femininus, Coris debueni, Pseudolabrus fuentesi* and *Thalassoma lutescens*) inhabiting Rapa Nui (Easter Island) in the southeastern Pacific Ocean, utilizing geometric morphometrics and aging methods. Age estimation based on otolith structure indicated that collected specimens were adults, ranging between 1 and 3 years old. Allometric analysis revealed low but significant variation (5.40%), primarily driven by changes in otolith width, resulting in distinct morphologies. Principal Component Analysis (PCA) of sagittal otoliths elucidated significant morphospace variation among species, with the first two PCs explaining 60.3% of the total variance. PC1 distinguished between elongated sagittal shapes (e.g., *P. fuentesi*) and robust sagittae (e.g., *T. lutescens*), while PC2 correlated with otolith roundness, delineating variations within species. Canonical variate analysis further highlighted differences in otolith shape, with significant variations detected among all species. The Discriminant function analysis showed high levels of discrimination accuracy for most of species’ pairs. Except for *C. debueni-P. fuentesi* (89%) and *C. debueni-T. lutescens* (96%), all other species pairs achieved 100% discrimination, highlighting the reliability of otolith shape as a distinguishing characteristic for the studied species. Overall, our findings emphasize the value of otolith shape analysis for characterizing and distinguishing Rapa Nui labrid species. This offers potential applications such as identifying prey within the digestive systems of large fish or bird predators in the vicinity of this isolated island in the South Pacific Ocean or archaeofauna studies.

## 1. Introduction

Otoliths play a key role for the hearing capabilities of fishes (Cruz and Lombarte, 2004; Deng et al., 2013). These structures, composed of calcium carbonate and structural proteins, grow by the deposition of concentric layers that are deposited sequentially, leaving an unequivocal record of the age and other life history traits from birth until the date of fish death (e.g., Panfili et al., 2002; Plaza et al., 2018). Furthermore, otoliths are characterized by having a highly species-specific shape, which is very useful for species and stock identification (Campana and Casselman, 1993; Moore et al., 2022). Additionally, otoliths are employed for ecomorphological studies (Victor, 1982; Zenteno et al., 2014; Tuset et al., 2018), spawning areas identification (e.g., Galley et al., 2006) and sex change determination, among other applications (Munday et al., 2009). Of all otolith types, the sagittal otolith exhibits the most remarkable feature, a species-specific morphology, with less variation within a species (Campana, 2004; He et al., 2018).

The shape of the otolith is influenced by its genotype, developmental stage, and the biotic and abiotic environments encountered during its lifetime (Mahé et al., 2019). Various methods exist for comparing otolith shape. Initially, linear measurements of distances were collected, and multivariate statistical methods were applied to analyze the shape of recording structures (i.e., traditional morphometrics) (Neilson et al., 1985; Takahashi et al., 2023). However, this approach has its limitations, such as the loss of information through shape simplification, and the high degree of multicollinearity between the measurements (Lishchenko and Jones, 2021). Another method is the elliptical Fourier analysis (EFA), which is a powerful method for describing outlines and extracting a significant percentage of shape variation that is easily visualized (Vieira et al., 2014). Variations in otolith shape can also be addressed using geometric morphometrics (GM), which is adaptation of multivariate statistics and graphics for studying phenotypic variation (Schaefer and Bookstein, 2009). To date, a number of studies have demonstrated GM is an efficient tool for distinguishing among species and visualizing modifications in otolith shape allometry and asymmetry (Ponton, 2006; Mahé et al., 2019; Cerda et al., 2021). Under the GM approach, otolith shape is defined by the coordinates of numerous points (landmarks) located along its perimeter (Palmer et al., 2010). These landmarks are transformed into a set of ordinary biometric variables, known as shape coordinates, which can then be individually regressed on the factors influencing them or the features of the systems they are presumed to affect (Schaefer and Bookstein, 2009).

In the family Labridae (wrasses, parrotfishes, and hogfishes), there is limited research on otolith morphology (but see Skeljo & Ferri 2012). This family comprises over 800 species, displaying a variety of body shapes, sizes, colors, and habitat preferences (Parenti & Randall, 2000; Parenti, 2011; Tea et al., 2024). Labrids inhabit diverse environments, including tropical shallow-water coral reefs, seagrass beds, and temperate rocky reefs. There is extensive evidence of niche partitioning and considerable trophic versatility among labrids. A given morphology may be associated with different feeding modes (Bellwood et al., 2006; Price et al., 2011), while some species show specialized feeding modes such as corallivory, planktivory, and molluscivory. Under such a great variability in feeding strategy and ecological niches seem reasonable infer that otolith shape can also vary markedly among these species, as suggested by ecomorphological otolith-based studies in other species (e.g., Assis et al., 2020; Bose et al., 2020).

Labrids are distributed across the Indo-Pacific Ocean, ranging from the warmer tropical waters to the colder waters of the southeastern Pacific, including oceanic islands of volcanic origin such as Rapa Nui (Easter Island). Located at the southeastern edge of Oceania, Rapa Nui lies approximately 2,800 km west of the Juan Fernández Islands off the coast of Chile, and 2,100 km east of the Pitcairn Islands (Randall and Cea, 2011). The island holds eleven labrid species representing four subfamilies and seven genera, including three small-range endemic species restricted to Rapa Nui and Motu Motiro Hiva (Salas y Gómez) (*Coris debueni, Novaculops koteamea*, and *Pseudolabrus semifasciatus*), three large-range endemic species (*Anampses femininus, Bodianus unimaculatus, Pseudolabrus fuentesi*) and five widespread species (*Anampses caeruleopunctatus, Cheilio inermis, Leptoscarus vaigiensis, Thalassoma lutescens* and *Thalassoma purpureum*) (Randall et al., 2005; Randall and Cea, 2011). Despite a few studies comparing insular species morphologically (Protti and Pequeño, 1999), research on otolith shape variability is absent.

This study aims to assess and compare the phenotypic variation in otolith shape among four labrid species collected in Rapa Nui waters, utilizing geometric morphometrics and aging methods. We show that the four species under study here have readily distinguishable otoliths. These results have interesting implications, as otoliths not only contribute to dietary analysis but also to the reconstruction of paleofauna (Weisler, 1993).

## 2. Materials and Methods

A total of 49 adult wrasses were collected in shallow waters of Hanga Roa, Rapa Nui (Easter Island), using scuba diving, between 2016 and 2018. The samples comprised a small-range endemic species, *C. debueni* (n=10); two large-range endemic species: *P. fuentesi* (n=9) and *A. femininus* (n=7), and one widespread species *T. lutescens* (n=23) (Table 1).

**Table 1.**
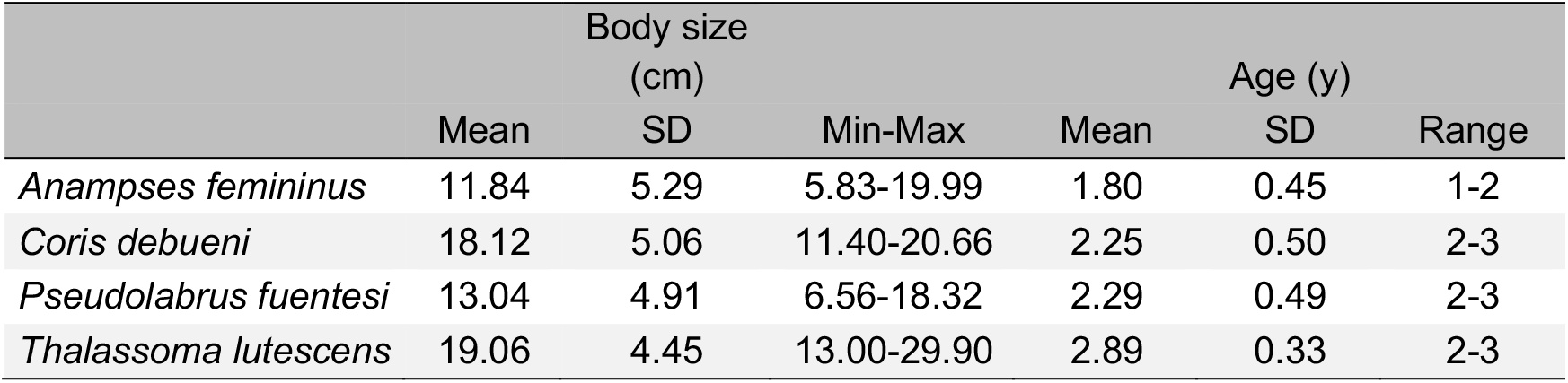
Body size (total length, cm) and age (years) ranges, mean and standard deviation (SD) of the wrasse species collected in Rapa Nui (Easter Island).

|  | Body size (cm) |  |  | Age (y) |  |  |
| --- | --- | --- | --- | --- | --- | --- |
|  | Mean | SD | Min-Max | Mean | SD | Range |
| *Anampses femininus* | 11.84 | 5.29 | 5.83-19.99 | 1.80 | 0.45 | 1-2 |
| *Coris debueni* | 18.12 | 5.06 | 11.40-20.66 | 2.25 | 0.50 | 2-3 |
| *Pseudolabrus fuentesi* | 13.04 | 4.91 | 6.56-18.32 | 2.29 | 0.49 | 2-3 |
| *Thalassoma lutescens* | 19.06 | 4.45 | 13.00-29.90 | 2.89 | 0.33 | 2-3 |

All specimens were measured in its total length (TL, cm). Left and right sagittal otoliths were extracted and mounted on a glass slide with Crystalbond™. Right otoliths were photographed with a replicate, under a stereomicroscope Leica EZ4E connected to a camera. The maximum Feret, from the anti-rostrum to the post-rostrum, and total area were measured. Then, a subset of otoliths was doble polished in sagittal position for age determination, using the MetaServ™ 250 device. A series of 800, 1000, and 2000 μm grit size of lapping film were used. For the estimation of the morphological variation of the otolith shape, 4 landmarks (LM) and 17 semi-landmarks (S-LM) were digitized into 130 photographs of the best-preserved right otoliths plus their replicates, using TPSDig 2.17. The landmark configuration represents the location of the rostrum (LM 1), anti-rostrum (LM 2), excisura minor (LM 9), and excisura major (LM 20). Semi-landmarks represent the distal border (S-LM 3-8), ventral border (S-LM 10-18), and the concave notch between LMs 1 and 2 (S-LM 19, 21) (Fig. 1). Semi-landmarks are useful to incorporate information about the curvatures (Ponton, 2006), and they were analyzed following Primost et al. (2015), utilizing TPSRelw 1.70. Shape information was extracted from the landmark coordinate with a generalized Procrustes analysis (GPA, Dryden and Mardia, 1998) using MorphoJ 1.5f (Klingenberg, 2011). These new coordinates were used for further statistical comparison. Centroid size (CS) was defined as the square root of the sum of the squared distances from the landmarks to the centroid and was used as a proxy for size. The estimation of the digitalizing error was obtained with a Procrustes ANOVA. To quantify the allometry, which reflects the covariation of traits among individuals, a multivariate regression of shape represented by Procrustes coordinates (landmarks coordinates after GPA) on size (log CS, Loy et al., 1998) was conducted. A principal component analysis was performed using the shape coordinates to identify the main axes of shape variation within and among species, as well as specific changes in the otolith contour. To compare otolith shapes among species, a Canonical Variate Analysis (CVA) based on the Mahalanobis distance was done. Significant differences were tested using a permutation test (10,000 rounds). Finally, a Discriminant Function Analysis was carried out to determine how discriminant may be the otolith shape for the identification of species, estimating the reliability of the discrimination using leave-one-out cross-validation test. All statistical analyses were run utilizing the software MorphoJ 1.5f.

**Fig. 1.**
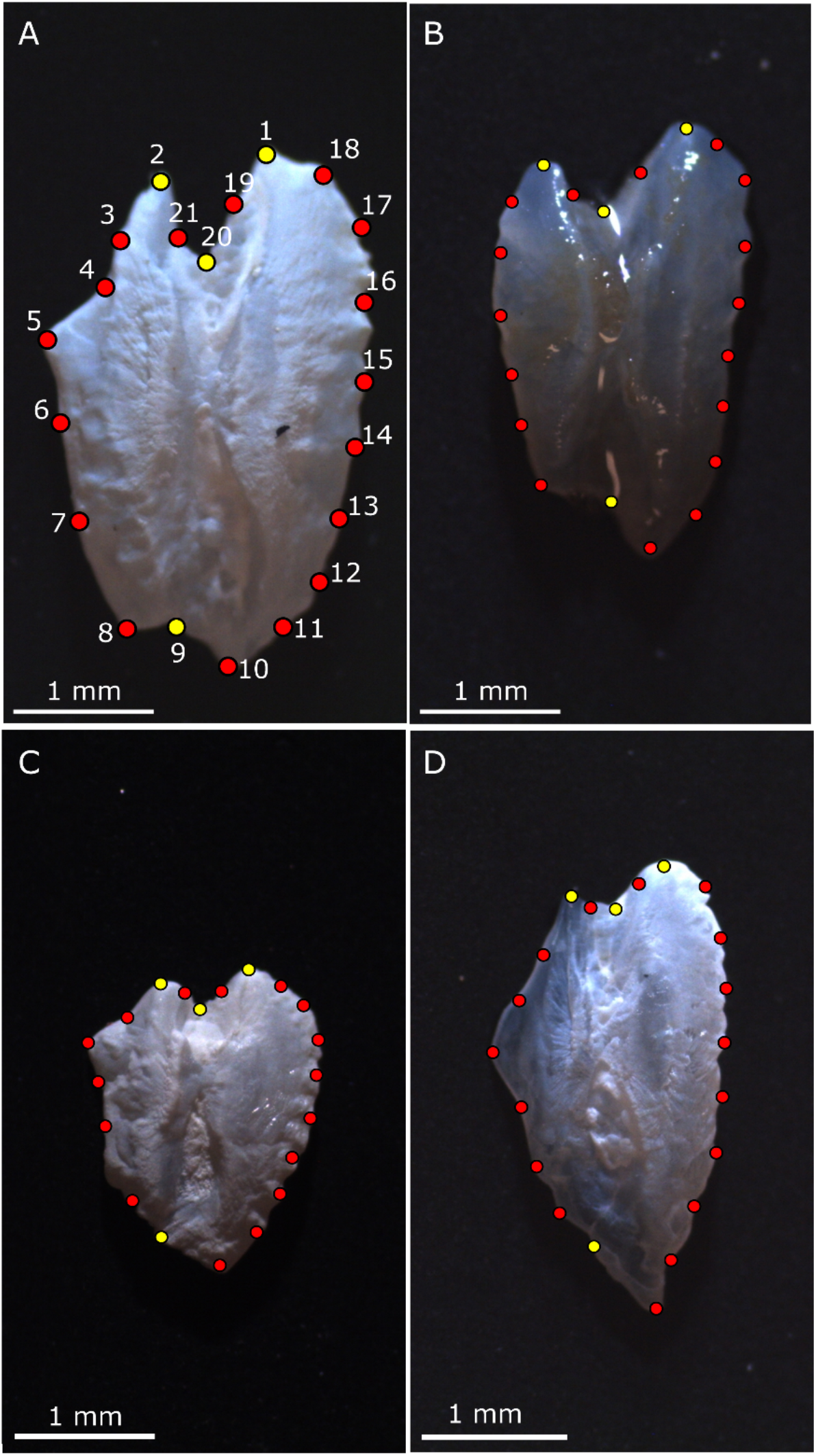
Otolith shape morphology of wrasses from Rapa Nui, indicating the location of landmarks (yellow dots) and semi-landmarks (red dots). A) *Thalassoma lutescens*, B) *Anampses femininus*, C) *Coris debueni* and D) *Pseudolabrus fuentesi*.

## 3. Results

The total length of specimens varied from 5.83 (*Anampses femininus*) to 29.90 cm (*Thalassoma lutescens*) (Table 1). The otolith size (i.e., maximum Feret) varied from 2.34 (*A. femininus*) to 4.50 mm (*Pseudolabrus fuentesi*). The age estimated by the otolith structure indicated that those species collected at Rapa Nui were adults (between 1 and 3 years, Table 1).

Measurement error, estimated by the Procrustes ANOVA, was negligible, since the individual mean square largely exceeded to the error, and accounted only 3.06% (Viscosi & Cardini, 2011). The allometry was low (5.40%), but significant (permutation test against the null hypothesis of independence, *P* = 0.023), represented by a change in the width of the otolith shape, i.e., starting with a wider dorsal border, and finishing with an enlargement of the otolith axis, particularly the rostrum, and a pointed anti-rostrum, and a narrowing of the notch (Fig. 2).

**Fig. 2.**
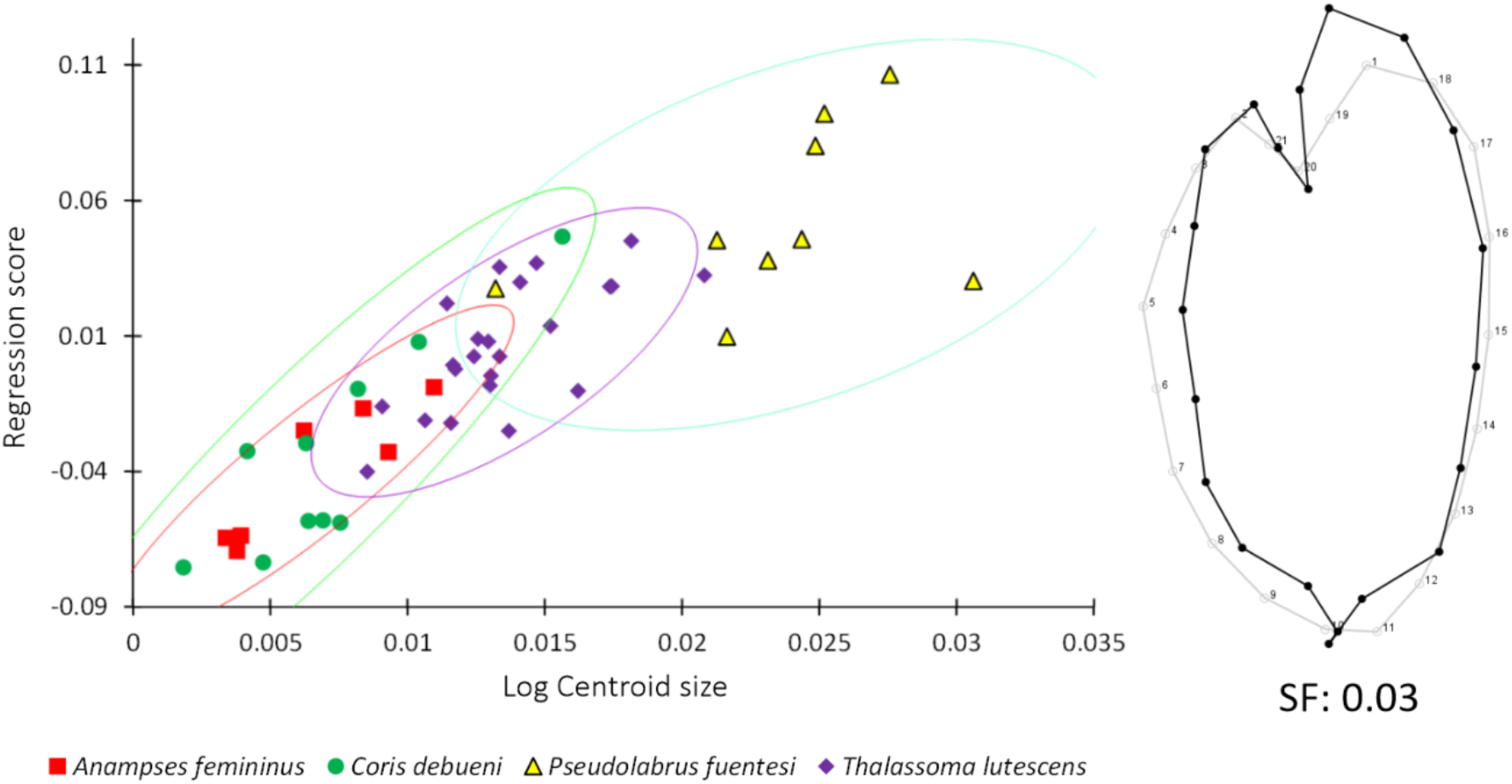
Multivariate regression between shape and log centroid size (i.e., trait size) representing the allometry in the otolith shape of wrasses from the southeastern Pacific Ocean. Gray (black) wireframes represent otolith shape changes related to otolith size. SF = scale factor; Confidence Ellipse: Probability 95%.

The morphospace of sagittal otoliths from wrasses of the southeastern Pacific Ocean was explained by 38 principal components (PCs), from which PC1 and PC2 explained 40.2 and 20.1% of the total variance, respectively (Fig. 3). The PC1 explained the differences between elongated sagittal shape with a pointed postrostrum and rostrum, and a small notch (*P. fuentesi*), to robust sagittae, with crenulations in the dorsal border with a deeper notch and similarly extended rostrum and antirostrum (*T. lutescens*) (Fig. 3). The PC2 was related to the roundness of the otoliths, ranging from otoliths with wider dorsal-ventral axis and similarly extended rostrum and antirostrum (*Coris* and *Anampses*), to otoliths with a narrower dorsal border, elongated rostrum and deeper notch (*T. lutescens*) (Fig. 3).

**Fig. 3.**
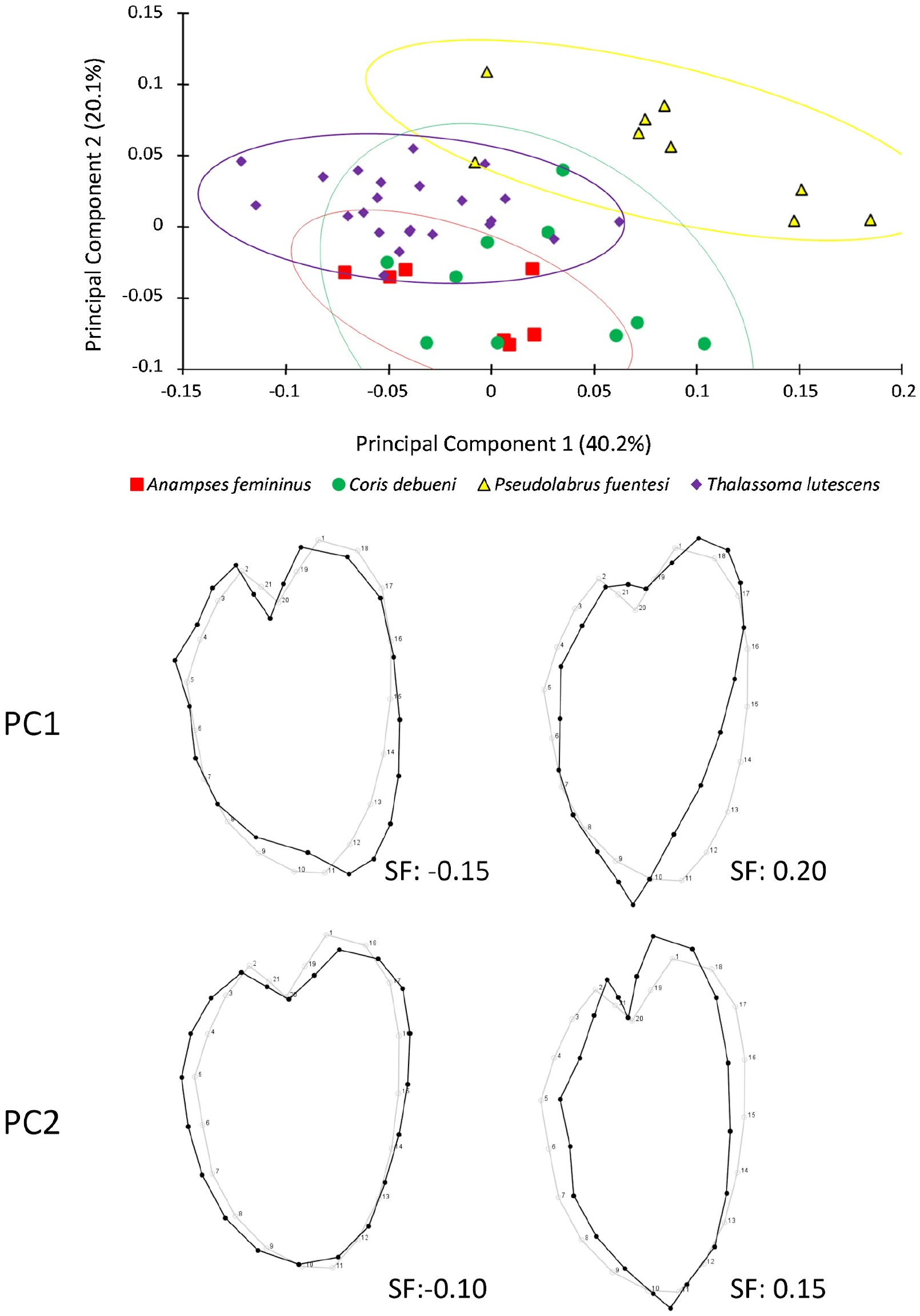
(Top) Principal component analysis (PCA) of the morphospace of sagittal otoliths of wrasses from oceanic islands from the southeastern Pacific Ocean. (Bottom) Grey wireframe represents the consensus shape, and black wireframes represent the target shape; numbers below the wireframes represent the scales in PC1 and PC2, respectively. SF: Scale factor; Confidence Ellipse: Probability 95%.

The comparison of otolith shape among species was explained by three canonical variates. CV1 delineated differences in the lengthening of the postrostrum of the sagitta, ranging from those with a pointed postrostrum (*Pseudolabrus*) to those with a rounded and wider posterior border (*Anampses*). CV2 explained variations in the depth of the notch, spanning from a shallow depression of the notch (*Coris*) to otoliths with a greater distance between the excisura major and the rostrum (*Thalassoma*, Fig. 4).

**Fig. 4.**
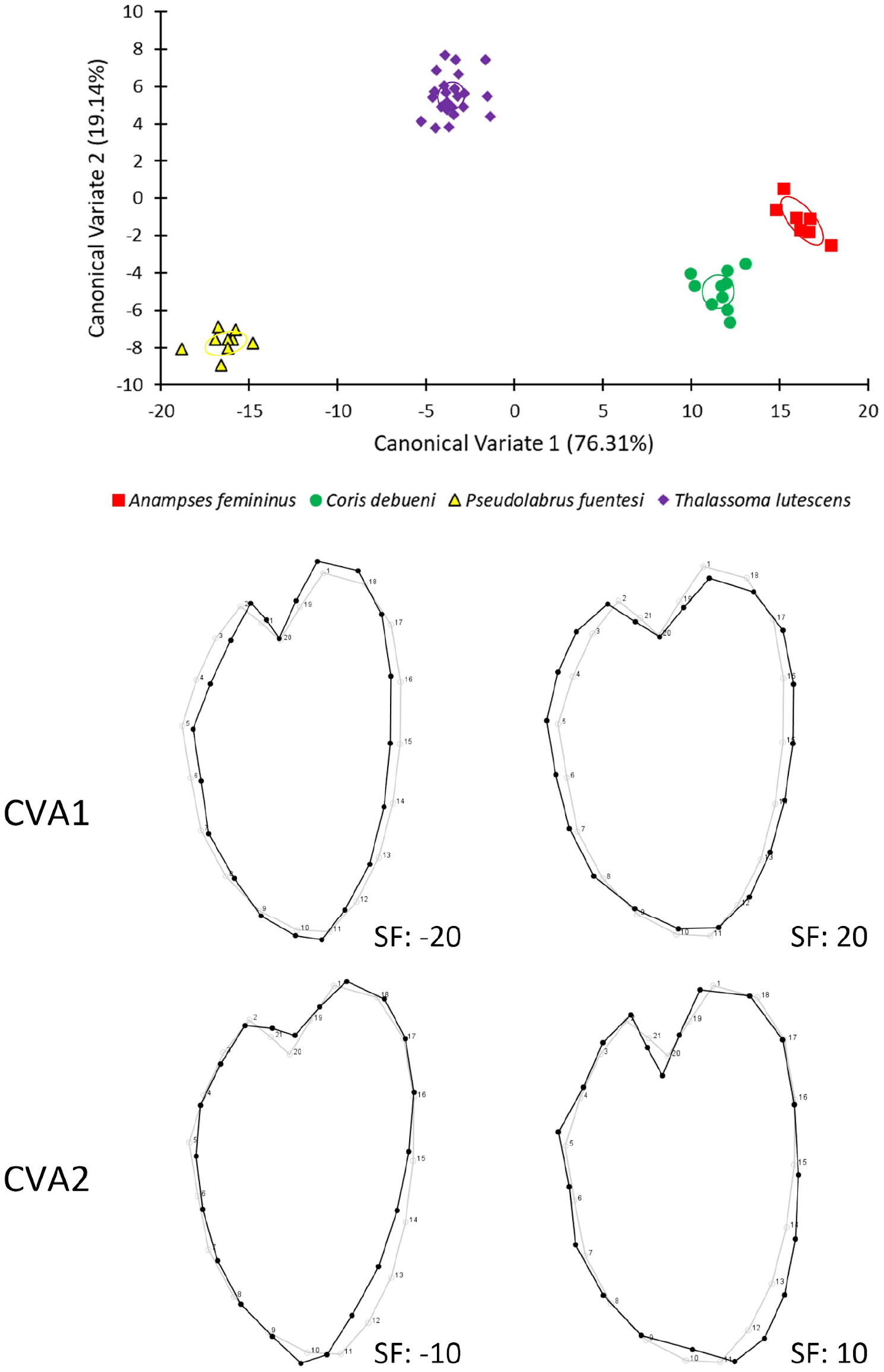
Canonical Variate Analysis (CVA) comparing otolith shape of wrasses from oceanic islands of the southeastern Pacific Ocean. SF = scale factor.

Shape comparison, based on the Mahalanobis distances, revealed significant differences among all studied species (permutation test, *P <* 0.001). The discriminant function analysis detected that most of the paired comparisons discriminated >90% correctly (Table 2). The least distinct pairs were *C. debueni-P. fuentesi* (cross-validation, 89% correct), and *C. debueni-T. lutescens* (cross-validation, 96%). However, in all other paired comparisons, both the discriminant function and the cross-validation indicated 100% discrimination among species. Therefore, the otolith shape of wrasses serves as a reliable discriminator for identifying four out of 10 species inhabiting Rapa Nui.

**Table 2.** Results of the Discriminant Function Analysis and the classification/misclassification for the leave-one-out cross-validation for the otolith shape between wrasse species from Rapa Nui.

| Comparison | Allocated from<br>Discriminant function | Allocated accuracy by<br>cross-validation | Procrustes Distance | *P* |
| --- | --- | --- | --- | --- |
| *Anampses-Coris* | 100 | 100 | 0.061 | 0.001 |
| *Anampses-Pseudolabrus* | 100 | 100 | 0.150 | 0.0001 |
| *Anampses-Thalassoma* | 100 | 100 | 0.089 | 0.0001 |
| *Coris-Pseudolabrus* | 100 | 86.8 | 0.120 | 0.0001 |
| *Coris-Thalassoma* | 100 | 96.8 | 0.092 | 0.0001 |
| *Pseudolabrus-Thalassoma* | 100 | 100 | 0.137 | 0.0001 |

## 4. Discussion

The analysis of sagittal otolith shape variability in four labrid species from Rapa Nui allowed for description and comparison at both intra- and interspecific levels. The main shape changes were observed between elongated sagittae shape with a pointed postrostrum and rostrum, and a small notch (*T. lutescens*), and robust sagittae with crenulations in the dorsal border, deeper notch and similarly extended rostrum and antirostrum (*P. fuentesi*). The second most relevant shape variation was related to the roundness of the otoliths, ranging from otoliths with a wider dorsal-ventral axis and similarly extended rostrum and antirostrum (*C. debueni* and *A. femininus*), to otoliths with a narrower dorsal border, elongated rostrum and deeper notch (*T. lutescens*). Therefore, the extension of the postrostrum and rostrum, and the depth of the notch represented good morphological characteristics for the discrimination of these species from Rapa Nui.

The family Labridae shows relatively smaller otoliths with large variability in shape (Cruz and Lombarte 2004), even within the same genus (e.g., *Symphodus*, Skeljo and Ferri 2012). Paxton (2000) suggested that the small size in coastal fish species may be associated with background noise and/or excessive movement of heavy otoliths in rough seas. Noticeable otolith shape differences were obeserved among species from Rapa Nui, showing a high degree of discrimination (>86%). The high interspecific variation in otolith morphology in closely related species found in the current study could suggests that shape is mainly under genetic control and can be used as natural tags for identifying and separating species, such as suggested by previous works (e,g,, Tuset et al., 2003; Skeljo and Ferri 2012). However, we cannot disentangle whether the differences in otolith shape found among species could also be driven by marked differential ecomorphological pattern, as recently demonstrated by Bose et al (2020) in four species of African cichlid fishes.

The otolith shape of *C. debueni* was similar to that of *C. julis*; both of them are cuneiform, and margins are entire to sinuate (Cruz and Lombarte, 2004; Skeljo and Ferri 2012); however, sagittae of *C. debueni* showed a more rounded shape compared to those from *C. julis*. The large and deep notch is observed also in sagittae from other labrids, such as *Symphodus tinca, S. cireneus*, and *Labrus merula* (Cruz and Lombarte, 2004; Skeljo and Ferri 2012). It is noticeably absent in the round otoliths of *Xyrichthys novacula* (Cruz and Lombarte, 2004), in juvenile *Pseudolabrus gayi* (Landaeta et al., 2022), and the triangle-like shape of *Pseudolabrus fuentesi* (this study).

The allometry, considering all four species, was low but significant, representing only 5.40% of the total variance. The interspecific allometric effect was mainly driven by a change in the width of the otolith shape, from a wider dorsal border to an enlargement of the otolith axis, with a pointed anti-rostrum, a larger rostrum, and a deeper notch. In the present case, all specimens used were adults of 1-3 yrs, and therefore, the ontogenetic allometry was non-significant at the intra-specific level, consistent with findings from other studies (e.g., Skeljo and Ferri 2012). Differences in otolith size and shape throughout ontogeny may arise due to variations in growth axis reshaped by environmental conditions experienced by the individuals in several micro-habitats from an island (slope vs estuary, Vignon, 2012). Therefore, the lack of (or reduced) intraspecific variability observed in sagittal otoliths of labrids from Rapa Nui may be due to the proximity of the sampling locations (nearby Hanga Roa), a similar density of fish in the sampled area (∼0.2 ind. m^-2^, Wieters et al., 2014), as well as the reduced variety of habitats (i.e., weak vertical zonation pattern, dominated by *Porites* corals, Wieters et al., 2014) compared to those from other islands of the Indo-Pacific (Vignon, 2012). It would be interesting to compare present findings with individuals collected from the southern coast of Rapa Nui, which is significantly more exposed.

In labrids, otoliths have been proven effective in determining various early life history traits, including size at hatch, pelagic larval duration (Victor 1986), recruitment size, and settlement patterns (Victor, 1982; Landaeta et al., 2022), as well as growth and mortality (Welsford, 2003; McBride and Richardson, 2007; Skeljo et al., 2012). Rapa Nui and Motu Motiro-Hiva are recognized as centers of endemism within the Indo-Pacific region. In this sense, it remains crucial to further investigate the macro- and microstructure of otoliths of endemic fishes from Easter Island to better understand their adaptations to life on this extremely isolated oceanic island.

## Availability of data and material

Raw coordinates of the geometric morphometric analysis are available as Supplementary Material.

## Code availability

Not applicable.

## Author’s contribution

**AC** extracted and mounted otoliths, digitized the landmarks, and performed data analysis, **EDT** and **PSA** designed and collected fish specimens, **PSA** and **MFL** obtained funds and designed the collection and methodology, **GP** performed otolith data analyses, **FM** assisted with the geometric morphometrics and multivariate statistical analysis, **MFL** wrote the first version of the manuscript; the whole team wrote the final version of the manuscript.

## Acknowledgments

We are grateful for the assistance provided in the otolith processing by Jorge E. Contreras (Instituto de Fomento Pesquero, Chile), Claudeth Asencio and Osneider Palomino (Pontificia Universidad Católica de Valparaíso, Chile). We thank Rebeca Tepano, Nina, Taveke Olivares Rapu, Liza Garrido Toleado (SERNAPESCA), and Ludovic Tuki (Mesa del Mar, Te Mau O Te Vaikava O Rapa Nui) for their kind and generous support. All applicable institutional guidelines for the care and use of animals were followed. Specimens were collected under permits No. 724, March 8th, 2016 and No. 1060, March 23th 2018, obtained from the Chilean Subsecretary of Fishing. The Universidad Austral de Chile Ethical Care Committee and Biosecurity Protocol approved our use and handling of animals.

## References

Assis, I.O., da Silva, V.E.L., Souto-Vieira, D., Lozano, A.P., Volpedo, A.V., Fabré, N.N., 2020. Ecomorphological patterns in otoliths of tropical fishes: assessing trophic groups and depth strata preference by shape. Environ. Biol. Fish. 103, 349–361. 10.1007/s10641-020-00961-0

Bose, A.P.H., Zimmermann, H., Winkler, G., Kaufmann, A., Strohmeier, T., Koblmüller, S., Sefc, K.M., 2020. Congruent geographic variation in saccular otolith shape across multiple species of African cichlids. Sci. Rep. 10, 12820. 10.1038/s41598-020-69701-9

Baliga, V.B., Law, C.J., 2016. Cleaners amongst wrasses: phylogenetics and evolutionary patterns of cleaning behavior within Labridae. Mol. Phylogenet. Evol. 94(A), 424–435. 10.1016/j.ympev.2015.09.006

Bellwood, D.R., Wainwright, P.C., Fulton, C.J., Hoey, A.S., 2006. Functional versatility supports coral reef biodiversity. Proc. Royal Soc. B 273, 101–107. 10.1098/rspb.2005.3276

Bernardi, G., Bucciarelli, G., Costagliola, D., Robertson, D.R., Heiser, J.B., 2004. Evolution of coral reef fish Thalassoma spp. (Labridae). 1. Molecular phylogeny and biogeography. Mar. Biol. 144, 369–375. 10.1007/s00227-003-1199-0

Campana, S.E., 2004. Photographic Atlas of Fish Otoliths of the Northwest Atlantic Ocean. Can. Spec. Publ. Fish. Aquat. Sci. 133, NCR Research Press, Ottawa.

Campana, S.E., Casselman, J.M., 1993. Stock discrimination using otolith shape analysis. Can. J. Fish. Aquat. Sci. 50, 1062–1083. 10.1139/f93-123

Castro, L.R., Landaeta, M.F., 2002. Patrones de distribución y acumulación larval en torno de las islas oceánicas: Isla de Pascua y Salas y Gomez. Cienc. Tecnol. Mar 25, 131–145.

Cerda, J.M., Palacios-Fuentes, P., Díaz-Santana-Iturros, M., Ojeda, F.P., 2021. Description and discrimination of sagittae otoliths of two sympatric labrisomid blennies Auchenionchus crinitus and Auchenionchus microcirrhis using morphometric analyses. J. Sea Res. 173, 102063. 10.1016/j.seares.2021.102063

Cruz, A., Lombarte, A., 2004. Otolith size and its relationship with colour patterns and sound production. J. Fish Biol. 65, 1512–1525. 10.1111/j.1095-8649.2004.00558.x

Delrieu-Trottin, E., Liggins, L., Trnski, T., Williams, J.T., Neglia, V., Rapu-Edmunds, C., Planes, S. Saenz-Agudelo, P., 2018. Evidence of cryptic species in the blenniid Cirripectes alboapicalis species complex, with zoogeographic implications for the South Pacific. Zookeys, (810), p.127.

Delrieu-Trottin, E., Brosseau-Acquaviva, L., Mona, S., Neglia, V., Giles, E.C., Rapu-Edmunds, C., Saenz-Agudelo, P., 2019. Understanding the origin of the most isolated endemic reef fish fauna of the Indo-Pacific: Coral reef fishes of Rapa Nui. J. Biogeogr. 46, 723–733. 10.1111/jbi.13531

Deng, X., Wagner, H.-J., Popper, A.N., 2013. Interspecific variations of inner ear structure in the deep-sea fish family Melamphaidae. Anat. Rec. 296, 1064–1082. 10.1002/ar.22703

Dryden, I. L., Mardia, K. V., 1998. Statistical analysis of Shape. John Wiley and Sons, Chichester.

Fernández-Cisternas, I., Majlis, J., Ávila-Thieme, M.I., Lamb, R.W., Pérez-Matus, A., 2021. Endemic species dominate reef fish interaction networks on two isolated oceanic islands. Coral Reefs 40, 1081–1095. 10.1007/s00338-021-02106-w

Galley, E. A., Wright, P. J., Gibb, F. B., 2006. Combined methods of otolith shape analysis improve identification of spawning areas of Atlantic cod. ICES J. Mar. Sci. 63, 1710–1717. 10.1016/j.icesjms.2006.06.014

He, T., Cheng, J., Qin, J., Li, Y., Gao, T., 2018. Comparative analysis of otolith morphology in three species of Scomber. Ichthyol. Res. 65, 192–201. 10.1007/s10228-017-0605-4

Klingenberg, C. P., 2011. MorphoJ: an integrated software package for geometric morphometrics. Mol. Ecol. Resour. 11, 353–357. 10.1111/j.1755-0998.2010.02924

Landaeta, M.F., Figueroa-González, Y., Moyano, G., Vera-Duarte, J., Pérez-Matus, A., Plaza, G., 2022. Mismatch between shape changes, early growth, and condition for a temperate reef fish from an oceanic island. Mar. Freshw. Res. 73, 624–635. 10.1071/MF21084

Lishchenko, F., Jones, J.B., 2021. Application of shape analyses to recording structures of marine organisms for stock discrimination and taxonomic purposes. Front. Mar. Sci. 8, 667183. 10.3389/fmars.2021.667183

Loy, A., Mariani, L., Bertelletti, M., Tunesi, L., 1998. Visualizing allometry: Geometric morphometrics in the study of shape changes in the early stages of the two-banded sea bream, Diplodus vulgaris (Perciformes, Sparidae). Journal of Morphology 237, 137–146. 10.1002/(SICI)1097-4687(199808)237:2<137::AID-JMOR5>3.0.CO;2-Z

Mahé, K., Ider, D., Massaro, A., Hamed, O., Jurado-Ruzafa, A., Goncalves, P., Anastasopoulou, A., Jadaud, A., Mytilineou, C., Elleboode, R., Ramdane, Z., Bacha, M., Amara, R., de Pontual, H., Ernande, B., 2019. Directional bilateral asymmetry in otolith morphology may affect fish stock discrimination based on otolith shape analysis. ICES Journal of Marine Science 76, 232–243. 10.1093/icesjms/fsy163

McBride, R.S., Richardson, A.K., 2007. Evidence of size-selective fishing mortality from an age and growth study of hogfish (Labridae: Lachmolaimus maximus), a hermaphroditic reef fish. Bull. Mar. Sci. 80, 401–417.

Moore, B.R., Parker, S.J., Pinkerton, M.H., 2022. Otolith shape as a tool for species identification of the grenadiers Macrourus caml and M. whitsoni. Fish. Res. 253, 106370. 10.1016/j.fishres.2022.106370

Munday, P.L., Ryen, C.A., McCormick, M.I., Walken, S.P.W., 2009. Growth acceleration, behaviour and otolith check marks associated with sex change in the wrasse Halichoeres miniatus. Coral Reefs 28, 623–634. 10.1007/s00338-009-0499-3

Neilson, J.D., Geen, G.H., Chan, B., 1985. Variability in dimensions of salmonid otolith otolith nuclei: implications for stock identification and microstructure interpretation. Fish. Bull. 83, 81–89.

Palmer, M., Linde, M., Morales-Nin, B., 2010. Disentangling fluctuating asymmetry from otolith shape. Mar. Ecol. Prog. Ser. 399, 261–272. 10.3354/meps08347

Parenti, P., 2011. Checklist of the species of the families Labridae and Scaridae: An update. Smith. Bull. 13, 29–44.

Parenti, P., Randall, J.E., 2000. An annotated checklist of the species of the labroid fish families Labridae and Scaridae. Ichthyol. Bull. 68, 1–97.

Paxton, J.R., 2000. Fish otoliths: do sizes correlate with taxonomic group, habitat and/or luminescence? Philos. Trans. R. Soc. Lond. B Biol. Sci. 355, 1299–1303. 10.1098/rstb.2000.0688

Plaza, G., Cerna, F., Landaeta, M.F., Hernández, A., Contreras, J.E., 2018. Daily growth patterns and age-at-recruitment of the anchoveta Engraulis ringens as indicated by a multi-annual analysis of otolith microstructure across developmental stages. J. Fish Biol. 93, 370–381. 10.1111/jfb.13773

Ponton, D., 2006. Is geometric morphometrics efficient for comparing otolith shape of different fish species? J. Morph. 267, 750–757. 10.1002/jmor.10439

Price, S.A., Holzman, R., Near, T.J., Wainwright, P.C., 2011. Coral reefs promote the evolution of morphological diversity and ecological novelty in labrid fishes. Ecol. Lett. 14, 462–469. 10.1111/j.1461-0248.2011.01617.x

Primost, M. A., Bigatti, G., Marquez, F., 2015. Shell shape as indicator of pollution in marine gastropods affected by imposex. Mar. Freshw. Res. 67, 1948–1954. 10.1071/MF15233

Protti, M., Pequeño, G., 1999. Diferenciación morfológica entre Pseudolabrus fuentesi (Regan, 1913) y Pseudolabrus gayi (Valenciennes, 1839) (Perciformes, Labridae) por métodos multivariados. Rev. Biol. Mar. Oceanogr. 34, 269–279.

Randall, J. E., Cea, A., 2011. Shore fishes of Easter Island. University of Hawaii Press.

Randall, J. E., Cea, A., Meléndez, R., 2005. Checklist of shore and epipelagic fishes of Easter Island, with twelve new records. Bol. Mus. Nac. Hist. Nat, Chile 54, 41–55.

Schaefer, K., Bookstein, F.L., 2009. Does geometric morphometrics serve the needs of plasticity research? J. Biosci. 34, 589–599. 10.1007/s12038-009-0076-5

Skeljo, F., Ferri, J., 2012. The use of otolith shape and morphometry for identification and size-estimation of five wrasse species in predator-prey studies. J. Appl. Ichthyol. 28, 524–530. 10.1111/j.1439-0426.2011.01925.x

Skeljo, F., Ferri, J., Brcic, J., Petric, M., Jardas, I., 2012. Age, growth and utility of otolith morphometrics as a predictor of age in the wrasse Coris julis (Labridae) from the eastern Adriatic Sea. Sci. Mar. 76, 587–595. 10.3989/scimar.03521.07G

Takahashi, M., Wakefield, C.B., Saunders, B.J., Fairclough, D.V., Harvey, E.S., Newman, S.J., 2023. Efficacy of otolith morphometry for rapid discrimination of cryptic fishes. Estuar. Coast. Shelf Sci. 295, 108516. 10.1016/j.ecss.2023.108516

Tea, Y-K., Zhou, Y., Ewart, K.M., Cheng, G., Kawasaki, K., DiBattista, J.D., Ho, S.Y.W., Lo, N., Fan, S. (2024). The spotted parrotfish genome provides insights into the evolution of a coral reef dietary specialist (Teleostei: Labridae: Scarini: Cetoscarus ocellatus). Ecol. Evol. 14, e11148. 10.1002/ece3.11148

Teresa, F. B., Sazima, C., Sazima, I., & Floeter, S. R. (2014). Predictive factors of species composition of follower fishes in nuclear-follower feeding associations: a snapshot study. Neotrop. Ichthyol., 12(4), 913–919. 10.1590/1982-0224-20140041

Tuset, V.M., Lombarte, A., González, J.A., Pertusa, J.F., Lorente, M.J., 2003. Comparative morphology of the sagittal otolith in Serranus spp. J. Fish Biol. 63, 1491–1504. 10.1111/j.1095-8649.2003.00262.x

Tuset, V.M., Olivar, M.P., Otero-Ferrer, J.L., López-Pérez, C., Hulley, P.A., Lombarte, A., 2018. Morpho-functional diversity in Diaphus spp. (Pisces: Myctophidae) from the central Atlantic Ocean: Ecological and evolutionary implications. Deep Sea Res. I 138, 46–59. 10.1016/j.dsr.2018.07.005

Victor, B.C., 1982. Daily otolith increments and recruitment in two coral-reef wrasses, Thalassoma bifasciatum and Halochoeres bivittatus. Mar. Biol. 71, 203–208.

Victor, B.C., 1986. Duration of the planktonic larval stage of one hundred species of Pacific and Atlantic wrasses (family Labridae). Mar. Biol. 90, 317–326. 10.1007/BF00428555

Vieira, A.R., Neves, A., Sequeira, V., Barros Paiva, R., Serrano Gordo, R., 2014. Otolith shape analysis as a tool for stock discrimination of forkbeard (Phycis phycis) in the Northeast Atlantic. Hydrobiologia 728, 103–110. 10.1007/s10750-014-1809-5

Vignon, M., 2012. Ontogenetic trajectories of otolith shape during shift in habitat use: Interaction between otolith growth and environment. J. Exp. Mar. Biol. Ecol. 420-421, 26–32. 10.1016/j.jembe.2012.03.021

Viscosi, V., Cardini, A., 2011. Leaf morphology, taxonomy and geometric morphometrics: a simplified protocol for beginners. PloS one, 6(10), e25630. 10.1371/journal.pone.0025630

Wei, X., Zhu, G., 2022. Shape and ontogenetic changes in otolith of the ocellated icefish (Chionodraco rastrospinosus) from the Bransfield Strait, Antarctic. Zool. 153, 126025. 10.1016/j.zool.2022.126025

Weisler, M.I., 1993. The importance of fish otoliths in Pacific island archaeofaunal analysis. N. Z. J. Archaeol. 15, 131–159.

Welsford, D.C., 2003. Interpretation of otolith microstructure in the early life history stages of two temperate reef wrasses (Labridae). Mar. Freshw. Res. 54, 69–75. 10.1071/MF02018

Westneat, M.W., Alfaro, M.E., 2005. Phylogenetic relationships and evolutionary history of the reef fish family Labridae. Mol. Phylogenet. Evol. 36, 370–390. 10.1016/j.ympev.2005.02.001

Wieters, E.A., Medrano, A., Pérez-Matus, A., 2014. Functional community structure of shallow hard bottom communities at Easter Island (Rapa Nui). Lat. Am. J. Aquat. Res. 42, 827–844. 10.3856/vol42-issue4-fulltext-10

Zenteno, J.I., Bustos, C.A., Landaeta, M.F., 2014. Larval growth, condition and fluctuating asymmetry in the otoliths of a mesopelagic fish in an area influenced by a large Patagonian glacier. Mar. Biol. Res. 10, 504–514. 10.1080/17451000.2013.831176

